# Low affinity integrin states have faster binding kinetics than the high affinity state

**DOI:** 10.1101/2021.07.26.453735

**Authors:** Jing Li, Jiabin Yan, Timothy A. Springer

## Abstract

Integrin conformational ensembles contain two low-affinity states, bent-closed and extended-closed, and an active, high-affinity, extended-open state. It is widely thought that integrins must be activated before they bind ligand; however, one model holds that activation follows ligand binding. As ligand-binding kinetics are not only rate limiting for cell adhesion but also have important implications for the mechanism of activation, we measure them here for integrins α4β1 and α5β1 and show that the low-affinity states bind substantially faster than the high-affinity state. On and off-rate measurements are similar for integrins on cell surfaces and ectodomain fragments. Although the extended-open conformation’s on-rate is ∼20-fold slower, its off-rate is ∼25,000-fold slower, resulting in a large affinity increase. The tighter ligand-binding pocket in the open state may slow its on-rate. These kinetic measurements, together with previous equilibrium measurements of integrin conformational state affinity and relative free energy on intact cells, are key to a definitive understanding of the mechanism of integrin activation.

## INTRODUCTION

Integrins are a family of adhesion receptors that mechanically integrate the intracellular and extracellular environments and facilitate cell migration. Their α and β-subunits associate noncovalently to form an extracellular ligand-binding head and then form multi-domain ‘legs’ that connect to single-pass transmembrane and cytoplasmic domains with binding sites for cytoskeletal adaptor or inhibitory proteins (Fig. 1A). Integrins populate a conformational ensemble with three overall conformational states: the low-affinity bent-closed (BC) and extended-closed (EC) conformations and the high-affinity extended-open (EO) conformation (Fig. 1A). The equilibrium between these conformational states is allosterically regulated by extracellular ligand binding, intracellular adaptor/inhibitor binding (Bouvard *et al*, 2013; Iwamoto & Calderwood, 2015) and tensile force applied by the actin cytoskeleton on the integrin β-subunit that is resisted by ligand embedded in the extracellular matrix or on cell surfaces (Kim *et al*, 2011; Legate & Fassler, 2009; Li & Springer, 2017; Nordenfelt *et al*, 2016; Park & Goda, 2016; Sun *et al*, 2016; Zhu *et al*, 2008) (Fig. 1A). The EO conformation has ∼1000-fold higher binding affinity for ligand than the two closed conformations and is the final competent state to mediate cell adhesion and migration (Li & Springer, 2018; Li *et al*, 2017; Schürpf & Springer, 2011). Many previous studies have emphasized the importance of force in regulating integrin adhesiveness (Alon & Dustin, 2007; Astrof *et al*, 2006; Li & Springer, 2017; Nordenfelt *et al*., 2016; Nordenfelt *et al*, 2017; Sun *et al*, 2019; Zhu *et al*., 2008). Recent measurements of the intrinsic ligand-binding affinity of each conformational state and the equilibria linking them enabled a thermodynamic comparison of integrin activation models (Li & Springer, 2017, 2018; Li *et al*., 2017). Remarkably, only the combination of adaptor binding and cytoskeletal force can activate integrins in an ultra-sensitive manner, with the switch between on and off occurring over a narrow range of signal input {Kuriyan, 2012 #24442}; the large increase in length between the bent and extended conformations (Fig. 1A) is indispensable for switch-like integrin activation.

**Figure 1.**
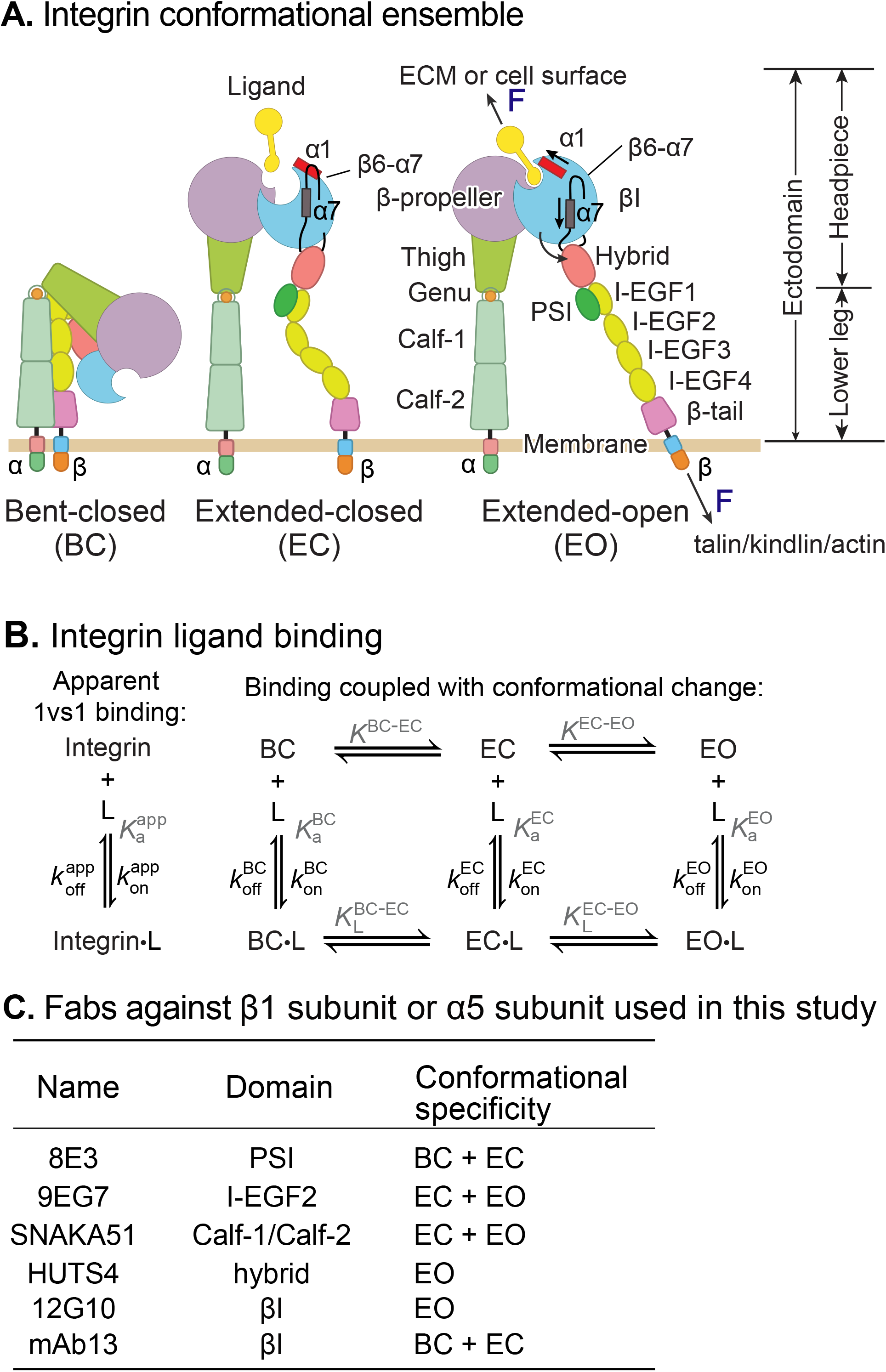
Ligand-interaction kinetics of integrin ensembles. **(A)** Three overall integrin conformational states (Luo *et al*, 2007). Individual domains are labeled next to the extended-open state. The structural motifs that move during opening (α1-helix, α7-helix and β6-α7 loop) are labeled in the βI domain of the EC and EO state. F represents tensile force exerted across ligand–integrin–adaptor complexes by the cytoskeleton and resisted by immobilized ligand. **(B)** Reaction scheme showing the apparent 1 vs. 1 kinetics of integrin and ligand binding (left), and the scheme required to correctly calculate ligand binding kinetics that takes into account the kinetics of conformational change (right). (**C**) Fabs utilized in this study, the integrin domains they bind, and their conformational specificity.

Despite these advances, thermodynamics cannot describe the sequence of events in a multi-step transition; furthermore, energy-driven processes such as cytoskeleton movements occur under non-equilibrium conditions. Ligand-binding on- and off-rates are key parameters that determine whether integrin encounter of ligand is timely and whether the ligand remains bound for a sufficiently long time for the integrin to exert its function in the presence of force.

Previous representative measurements (Dong *et al*, 2018; Kokkoli *et al*, 2004; Mould *et al*, 2014; Takagi *et al*, 2003) on integrin interaction with ligand have yielded kinetics on mixtures of conformational states, i.e., apparent on-and off-rates averaged over conformational states (Fig. 1B left). However, the ligand-binding kinetics of individual integrin conformational states remain unknown. These kinetics must be determined before we can understand how integrin function is regulated and how integrins work in concert with the cytoskeleton to provide traction for cell migration and firm adhesion for tissue integrity (Fig. 1B right).

Putting the question another way, what is the first step in inside-out integrin activation? In one view, talin binding inside the cell activates the integrin, presumably to the high affinity state, which then binds ligand. In another view, the first step is activation of the actin cytoskeleton, followed by binding of the integrin to ligand embedded in the extracellular environment and to talin incorporated in the actin cytoskeleton, which enables actin retrograde flow to elongate the lifetime of the high affinity integrin state.

For two classes of force-regulated adhesion molecules, each of which have a single low-affinity state and a single high-affinity state, selectins (Phan *et al*, 2006) and FimH (Yakovenko, 2015), the low-affinity conformation has a faster on-rate for ligand than the high-affinity conformation. If subsequent conformational change to the high affinity state is rapid, fast ligand binding kinetics to the low-affinity state efficiently couples ligand binding to stabilization by applied force of the high-affinity state, which has a long lifetime (Yakovenko, 2015). Work from our group on integrin αVβ6 showed that removal of the hybrid domain in the αVβ6 head resulted in a 50-fold increase in affinity for ligand yet decreased the apparent on-rate of ligand binding (Dong *et al*., 2018) suggesting that the open conformation has a lower on-rate than the BC and EC states. However, the intrinsic ligand-binding kinetics for each state of integrin αVβ6 could not be determined due to the lack of tools to stabilize specific conformational states.

In this study, we utilized well-characterized conformation-specific Fabs (Li & Springer, 2018; Li *et al*., 2017; Su *et al*, 2016) (Fig. 1C) to stabilize integrins α4β1 and α5β1 into defined ensembles containing only one or two of the three integrin conformational states and measured the ligand-binding kinetics of each defined ensemble. Together with previously determined intrinsic ligand-binding affinities and populations of conformational states (Li & Springer, 2018; Li *et al*., 2017), our measurements enable us to define ligand-binding kinetics intrinsic to each conformational state. For each integrin, the two closed states have indistinguishable on- and off-rates for soluble peptide and macromolecular fragment ligands. Remarkably, the on-rate for ligand of the low-affinity closed integrin conformations is ∼40-fold (α4β1) or ∼5-fold (α5β1) higher than for the high-affinity EO conformation. The ∼1,000-fold higher affinity of the EO conformation than the closed conformation is achieved by the ∼25,000-fold lower off-rate of the EO conformation for both α4β1 and α5β1 integrins. These findings show for two representative β1 integrins that most ligand binding occurs to the bent-closed and/or extended-closed states, followed by conformational change to the extended-open state. The rapidity of ligand binding measured here, if coupled with similarly rapid binding of actin cytoskeleton adaptors to integrins and conformational change among integrin states, could enable coincidence of these binding events, together with tensile force transmission if the ligand is embedded in an extracellular environment, to regulate integrin activation.

## RESULTS

### Ligand-binding kinetics of intact α4β1 and α5β1 on cell surfaces

We measured binding kinetics of intact α4β1 on Jurkat cells to two fluorescently labeled ligands, a phenylureide derivative of Leu-Asp-Val-Pro (FITC-LDVP) and a fragment of vascular cell adhesion molecule (VCAM) containing its first two domains (Alexa488-VCAM D1D2) (Fig. 2). Before adding ligands, cells were equilibrated with saturating concentrations of Fabs for 30 min at 22°C to stabilize specific conformational states (Li & Springer, 2018). Integrin extension, i.e. the EC and EO states, was stabilized with 4 µM 9EG7 Fab, which binds to the β1-subunit knee (Fig. 2B). The EO conformation was stabilized with a combination of 4 µM 9EG7 Fab and 2 µM HUTS4 Fab; the latter binds to the interface between the βI and hybrid domains and stabilizes the EO conformation (Fig. 2C). Ligand binding kinetics was monitored as mean fluorescence intensity (MFI) by flow cytometry without washing (Fig. 2). Beginning at about 10 minutes, a 500-fold higher concentration of unlabeled ligand was added to measure the kinetics of dissociation. Background MFI at each fluorescent ligand concentration, measured under identical conditions except in presence of 10 mM EDTA (Fig. S1), showed no significant difference at different time points during the association and dissociation measurements and was averaged across different time points and subtracted to obtain specific binding.

**Figure 2.**
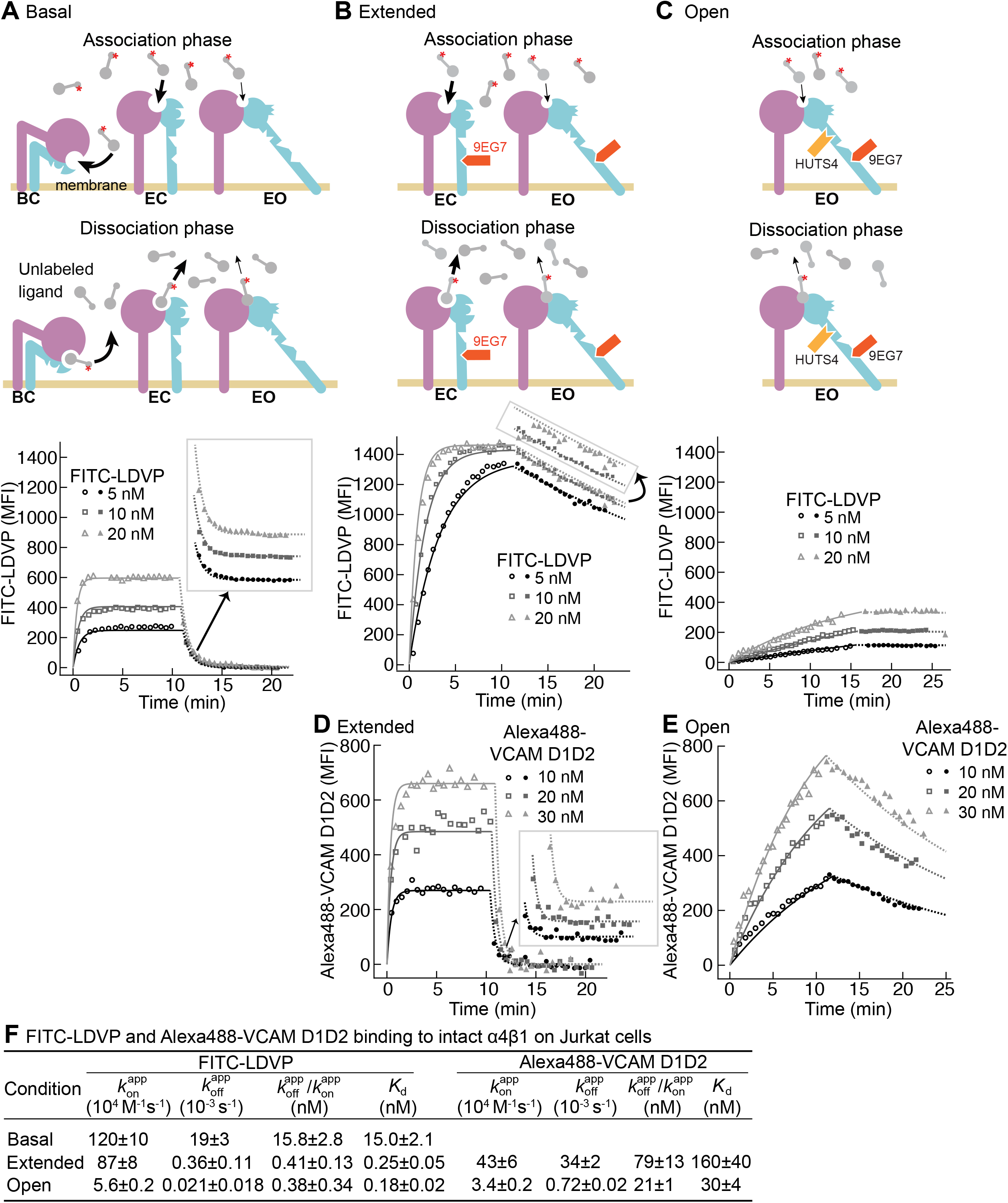
Binding kinetics of ligands to intact α4β1 on Jurkat cells. (**A-E**) Binding and dissociation of FITC-LDVP (A-C) and Alexa488-VCAM D1D2 (D-E) to α4β1 on Jurkat cells measured by flow cytometry. Cartoons in panel A, B, and C show the schemes for measuring ligand binding and dissociation in the association phase and dissociation phase in basal ensemble (A), extended ensembles (EC+EO states) stabilized with Fab 9EG7 (4 μM) (B and D), and open ensemble (EO state) stabilized with Fabs 9EG7 (4 μM) and HUTS4 (2 μM) (C and E), respectively. Specific MFI with the MFI in EDTA (Fig. S1) subtracted is shown as open (association) or filled (dissociation) symbols; fits are shown as thin lines as indicated in keys. **(F)** Tabulation of results. 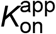 and 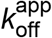 are from global fits of data at all ligand concentrations. 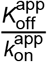 is also shown and compared to previous equilibrium *K*_d_ measurements (Li & Springer, 2018), except for Alexa488-VCAM D1D2 binding to extended states, which was measured here (Fig. S2). Errors for 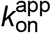 and 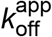 values are s.e. from global nonlinear least square fitting; errors for 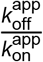 are propagated from errors of 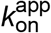 and 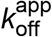 errors for *K*_d_ values are s.d. from three independent experiments.

Under basal conditions, with all three integrin states present in the ensemble, binding of FITC-LDVP to Jurkat cells reached equilibrium within 3 min (Fig. 2A). Upon addition of a 500-fold excess of LDVP, dissociation of FITC-LDVP was rapid and was 99.7% complete by 5 min (Fig. 2A). In contrast, both binding and dissociation of FITC-LDVP were slower when only the extended conformations (EC and EO) were present on Jurkat cells (Fig. 2B). Reaching steady state required ∼5 min after addition of 20 nM FITC-LDVP, ∼10 min with 10 nM ligand, and was not reached after 10 min with 5 nM ligand. After 10 min of dissociation, only 19.4% of ligand had dissociated (Fig. 2B). Association and dissociation were even slower when only the EO conformation was present (Fig. 2C). After 15 min of association, much less ligand had bound (Fig. 2C) than when both EC and EO conformations were present (Fig. 2B). Dissociation was also slower, with only 1.2% of bound ligand dissociating after 10 min (Fig. 2C).

VCAM D1D2 binds with ∼100-fold lower affinity than LDVP to α4β1 (Li & Springer, 2018). As a result, binding to the basal ensemble was too low to measure over the noise from unbound ligand; however, we were able to measure binding kinetics to intact α4β1 stabilized in the extended (EC+EO) and EO states (Fig. 2D and E). When the two extended conformations (EC and EO) were present, binding of all three concentrations of Alexa488-VCAM D1D2 (10nM, 20nM and 30nM) reached equilibria within 2 min. Upon addition of a large excess of LDVP, dissociation of Alexa488-VCAM D1D2 was also fast; 100% dissociated by 5min (Fig. 2D). Association and dissociation both became markedly slower when only the EO conformation of α4β1 was present (Fig. 2E).

To address the generality of these results, we studied another integrin and cell type by measuring binding of a fluorescently-labeled two-domain fragment of fibronectin (Alexa488-Fn3_9-10_) to intact α5β1 integrin on K562 cells (Fig. 3). The BC conformation of α5β1 integrin on K562 cells) is more stable than that of α4β1 integrin on Jurkat cells (Li & Springer, 2018). Therefore, to assure that the extended states (EC+EO) were saturably populated, they were stabilized with a combination of two Fabs, 6 µM 9EG7 Fab and 2 µM SNAKA51 Fab (Fig. 3A left). The EO state of α5β1 (Fig. 3B left) was stabilized with the same combination of Fabs as used for α4β1. Although binding affinity was too low to measure kinetics of the basal ensemble (Li *et al*., 2017), we were able to measure Alexa488-Fn3_9-10_ kinetics with the EC+EO and EO ensembles of intact α5β1 (Fig. 3). When α5β1 was stabilized in the EO conformation, Alexa488-Fn3_9-10_ bound and dissociated significantly more slowly than when both the EC and EO states of α5β1 were present in the ensemble (Fig. 3A and B). Faster binding and dissociation of Alexa488-Fn3_9-10_ from the EC+EO ensemble than EO showed that the EC state of α5β1 binds and dissociates faster than the EO state, just as found for α4β1.

**Figure 3.**
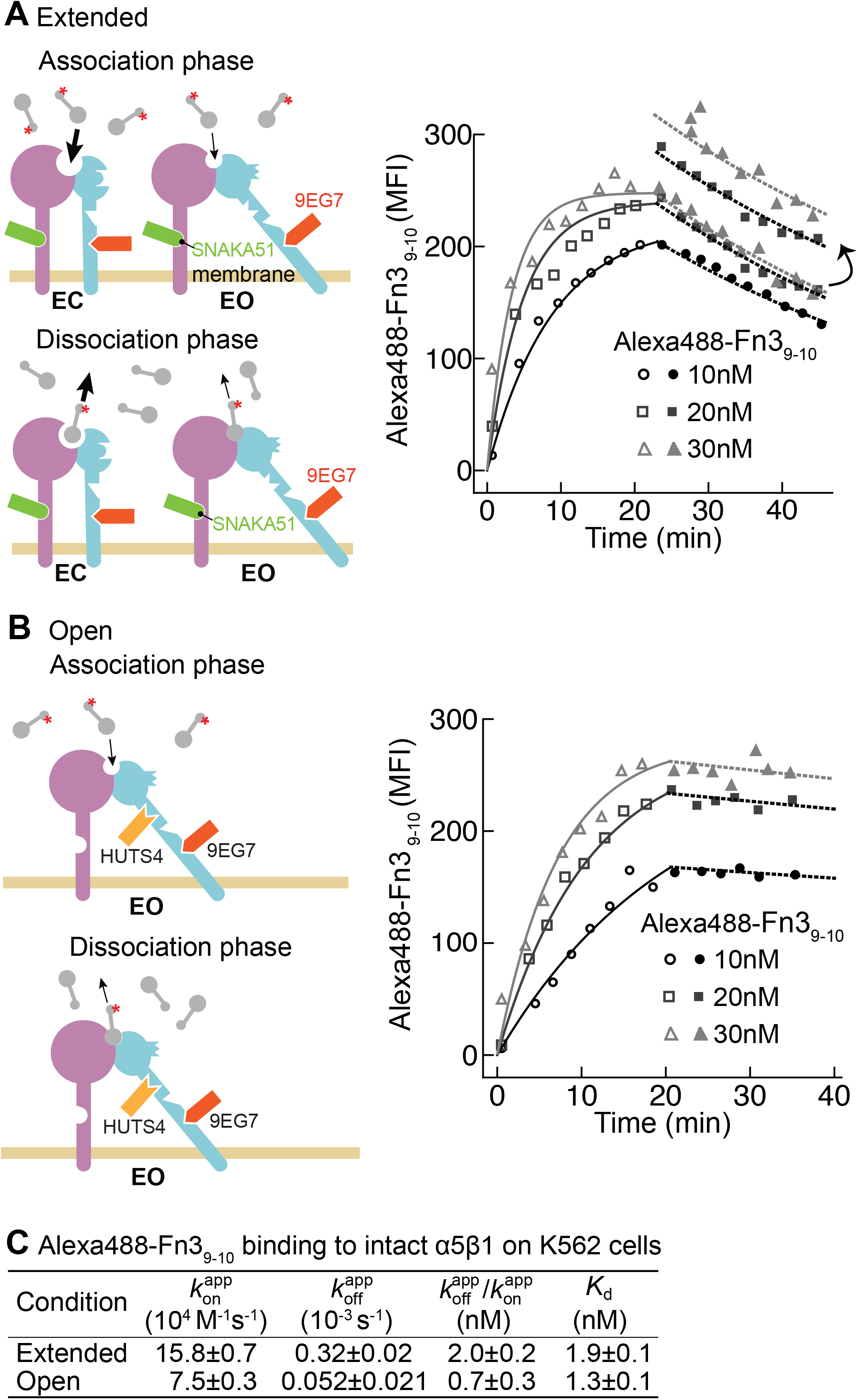
Binding kinetics of Alexa488-Fn3_9-10_ to α5β1 on K562 cells. **(A-B)** Binding of Alexa488-Fn3_9-10_ to α5β1 on K562 cells measured by flow cytometry. Measurements were on integrins in extended ensembles (EC+EO states) in presence of Fabs 9EG7 (6 μM) and SNAKA51 (2 μM) (A) or in the open (EO state) in presence of Fabs 9EG7 (6 μM) and HUTS4 (2 μM) (B), as illustrated in the cartoons. MFI with background in EDTA subtracted (Fig. S1) is shown as symbols and fits are shown as lines as explained in keys; the association phase has open symbols and solid lines, and the dissociation phase has filled symbols and dashed lines. (**C)** Tabulation of results. 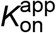 and 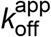 are from global fits and errors are from non-linear least square fits. 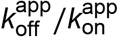 is also shown with propagated error and compared to previous equilibrium *K*_d_ measurements (Li *et al*., 2017).

To quantify the binding kinetics of intact α4β1 and α5β1 under each condition, we globally fit the traces of specific binding in both association and dissociation phases at each concentration of fluorescently labeled ligand to the 1 vs. 1 Langmuir binding model to determine the apparent on- and off-rates, 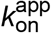 and 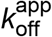 (Fig. 2F and Fig. 3C). The ratio of the apparent off- and on-rates, 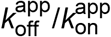, agrees reasonably well with the equilibrium dissociation constant, *K*_d_, previously determined by saturation binding (Figs. 2F and 3C) (Li & Springer, 2018; Li *et al*., 2017). These agreements suggest that the 1 vs. 1 Langmuir binding model can reasonably fit the kinetic data. Overall, these results show that ligand binds to and dissociates from the EO conformation more slowly than from the BC and EC conformations. The kinetics measured here for the basal and EC+EO ensembles are apparent, because they include contributions from distinct conformational states present in these ensembles. In contrast, EO state kinetics are measured exactly because EO is the only state present in the EO ensemble. In the final section of Results, we will use previous measurements of the populations of the states in each ensemble to calculate the on- and off-rates for conformations within mixtures of states.

### Binding kinetics of soluble α5β1 ectodomain for Fn3_9-10_

We utilized bio-layer interferometry (BLI) (Wallner *et al*, 2013) to measure the kinetics of binding of an ectodomain fragment of α5β1 to the biotin-labeled Fn3_9-10_ fragment of fibronectin immobilized on streptavidin biosensors (Fig. 4). The ectodomain was truncated just prior to the transmembrane domains of the α5 and β1 subunits and was expressed in a cell line containing a glycan processing mutation so that it had high-mannose rather than complex-type N-glycans. Truncation of α5β1 and high mannose glycoforms raise the free energy of the BC conformation relative to the EO conformation, so that the population of the EO state in the basal ensemble increased from 0.11% in intact α5β1 to 4.6% in the high-mannose ectodomain fragment (Li *et al*., 2017). The practical consequence of the increase in population of the EO state in the α5β1 ectodomain basal ensemble was that it raised basal ectodomain ensemble affinity and, in contrast to intact α5β1 on K562 cells, enabled us to measure basal ensemble Fn3_9-10_ binding kinetics (Fig. 4A).

**Figure 4.**
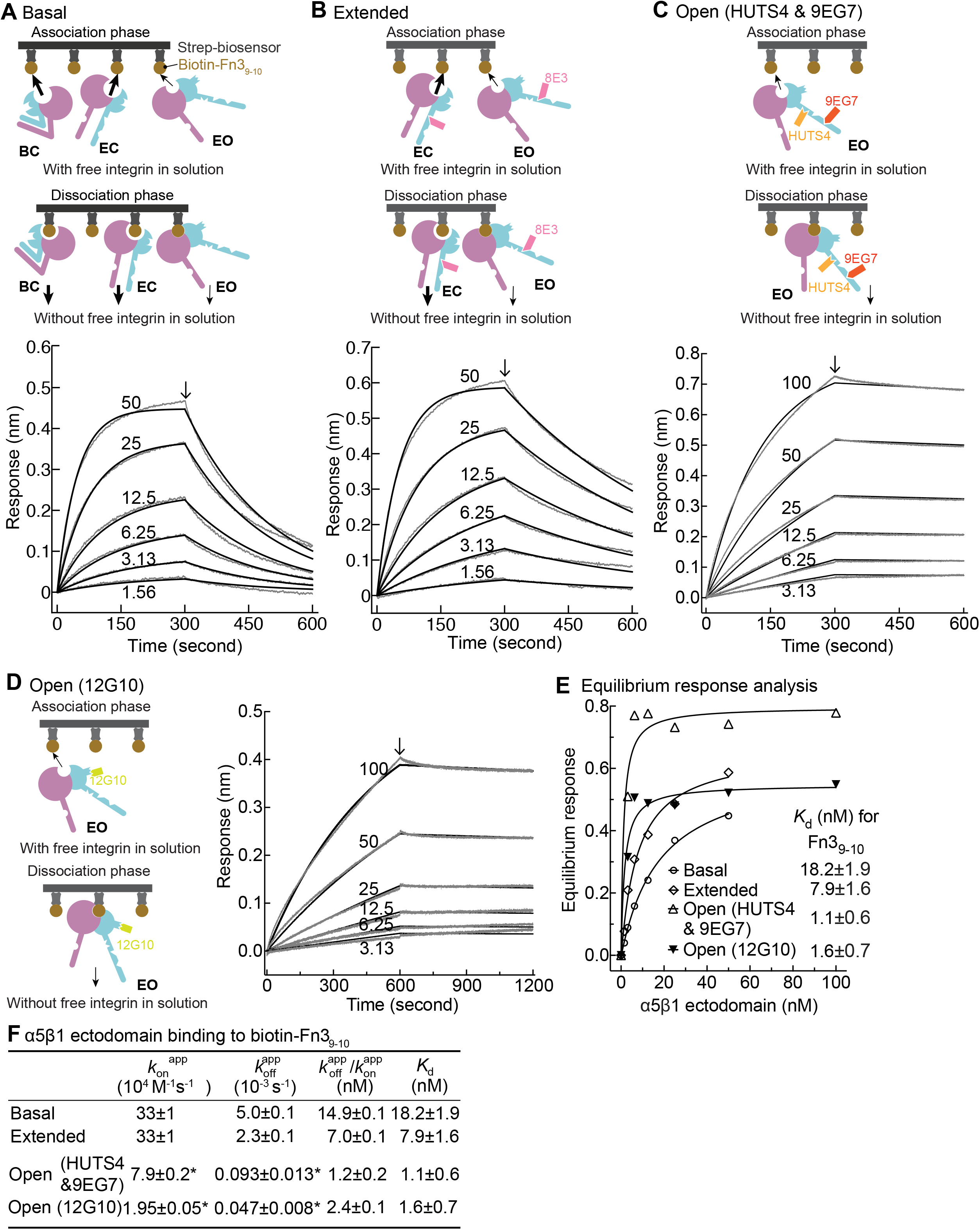
Binding kinetics of α5β1 ectodomain to Fn3_9-10_. **(A-D)** Binding of unclasped high-mannose α5β1 ectodomain measured with BLI. Schemes for measuring ligand binding and dissociation in the association phase and dissociation phase are shown in each panels’ cartoon. α5β1 ectodomain (analyte) at the indicated concentrations in nM was bound to biotin-Fn3_9-10_ immobilized on streptavidin biosensors without Fab (**A**) or with 2 μM Fab 8E3 (**B**), or with 2 μM 9EG7 and 5 μM HUTS4 Fabs, (**C**) or with 1 μM Fab 12G10 (**D**). Arrows mark the start of the dissociation phase. Response curves are in gray and fitting curves in black. (**E**) The equilibrium binding (response) was calculated from 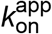 and 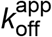 values at each α5β1 ectodomain concentration and fit to a dose response curve to calculate *K*_d_ values. These values serve as a check on the 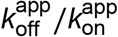 values in F. (**F)** Tabulation of *K*_d_ values from equilibrium response analysis in Panel E, 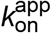 and 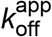 values from nonlinear least square fit of data in Panel A-D with 1 vs. 1 Langmuir binding model, and 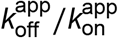. Errors without ^*^ for 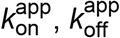 and *K*_d_ are fitting errors from nonlinear least square fits; errors for 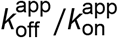 are propagated from errors of 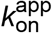 and 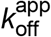. Errors with ^*^ are difference from the mean of two independent measurements.

Binding kinetics were measured by transferring Fn3_9-10_ biosensors to wells containing the α5β1 ectodomain in the absence or presence of conformation-stabilizing Fabs. Dissociation kinetics were measured by transfer of sensors to wells lacking the integrin but containing identical Fab concentrations (Fig. 4 A-D cartoons). Equilibrium *K*_d_ values were previously shown to be independent of the Fab used to stabilize a particular state (Li *et al*., 2017). However, we were concerned that binding of Fabs, particularly those that bind close to ligand binding sites, might slow kinetics and therefore tested this by varying the Fabs used to stabilize the EO state.

The kinetic curves showed that the α5β1 ectodomain EO state associated more slowly than the mixtures with the closed states and also dissociated more slowly (Fig. 4A-D) as confirmed in the tabulated results (Fig. 4F). Overall, these differences among ensembles resembled those found for the EC+EO ensemble and EO state of intact α5β1 on K562 cells and extended measurements to the basal α5β1 ensemble. The on and off-rates of the EO state for Fn3_9-10_ determined in the presence of 12G10 Fab were 4-fold and 2-fold lower, respectively, than those determined in the presence of 9EG7&HUTS4 Fabs (Fig. 4C, 4D and 4F). As 12G10 Fab binds close to the ligand-binding site in the β1 domain (Fig. 1A), we use 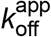 and 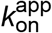 kinetics determined with the 9EG7, 8E3, SNAKA51 & HUTS4 Fabs, which bind far from the ligand-binding site, for calculating true (*k*_off_ *k*_on_ kinetic rates for each state in the final section of Results.

### The off-rate of the closed states

Due to the low affinities of the closed states there was too little binding to directly measure *k*_on_ or *k*_off_ in presence of saturating closure-stabilizing Fabs. We therefore used another approach. We first allowed ligand binding to integrins to reach steady state in the absence of a closure-stabilizing Fab. We then added different concentrations of closure-stabilizing Fab mAb13 and measured dissociation kinetics (Figs. 5-6). Dissociation of the ligand from the EO state is very slow as shown above and is negligible in our experimental time scale. At high Fab mAb13 concentrations, when the EO ligand-bound state (EO•L) converts to either BC•L or EC•L (they are grouped together here as (C•L), mAb13 Fab binds and prevents back-conversion to EO•L (Fig. 5A, B). After saturating concentrations of Fab mAb13 are added to basal or EO+EC ensembles pre-equilibrated with ligand, the effective off-rate is contributed by two steps, the conformational change from EO•L to C•L and the dissociation of ligand from mAb13-bound C•L (mAb13•C•L) (Fig.5A, B). Thus, the observed off-rate at saturating concentration of mAb13 Fab is contributed by the rates of both steps and permits the determination of the lower limit of 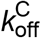.

**Figure 5.**
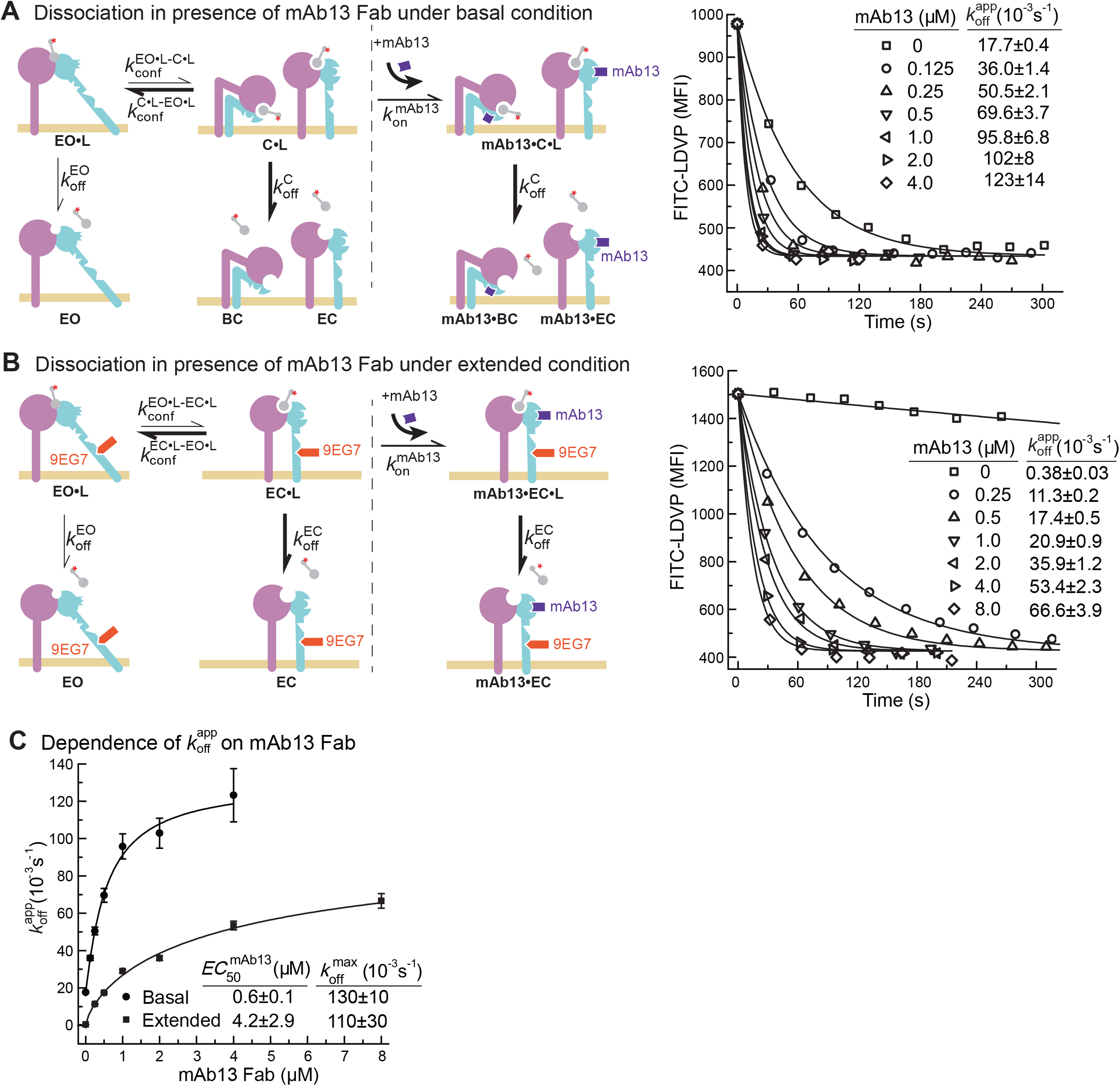
Dissociation of FITC-LDVP from α4β1 on Jurkat cells in presence of closure-stabilizing Fab. (**A-B)** FITC-LDVP dissociation from basal or extended ensembles of intact α4β1 on Jurkat cells measured using flow cytometry. FITC-LDVP (20nM) was incubated with Jurkat cells in absence (A) or in presence of extension-stabilizing Fab 9EG7 (4 μM) (B) for 10 minutes to reach steady state. Then, 10 μM unlabeled LDVP together with indicated concentrations of mAb13 Fab were added. Observed MFI (*MFI*_obs_) values as a function of time at indicated mAb13 Fab concentrations were globally fitted to 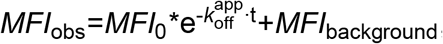, with *MFI* at the start of dissociation (*MFI*_0_) and background *MFI* (*MFI*_background_) as shared parameters and 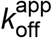 as the individual fitting parameter at each mAb13 Fab concentration. (**C**) Dependence of 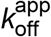 on mAb13 Fab concentration. 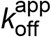 at each mAb13 Fab concentration in panels A and B were fitted to dose response curves to determine the maximum off-rate at saturating mAb13 Fab concentration, 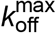, and the mAb13 Fab concentration when the off-rate reaches half of the maximum, 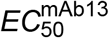. All errors are from nonlinear least square fits.

We measured FITC-LDVP dissociation from basal or extended ensembles of α4β1 on Jurkat cells after addition of a range of mAb13 Fab concentrations (Fig. 5A-B). Saturable binding of mAb13 Fab to nascent cell surface C•L was evident from the approach to a plateau of 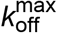 (Fig. 5A-C). The 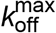 values measured for LDVP dissociation from basal and extended α4β1 ensembles on Jurkat cells were similar and within error of one another, with an average of ∼120 ^*^10^−3^ /s (Fig. 5C).

Similarly, we measured Fn3_9-10_ dissociation from basal or extended ensembles of the α_5_β_1_ ectodomain (Fig. 6). The effect of mAb13 Fab on increasing *k*_off_ was saturable, as shown by approach to a plateau (Fig.6A-C). The fit to a saturation dose response curve yielded 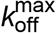 values for the basal and extended ensembles of (1600 ± 100) ^*^10^−3^ /s and (1900 ± 100) ^*^10^−3^/s, respectively (Fig. 6C).

**Figure 6.**
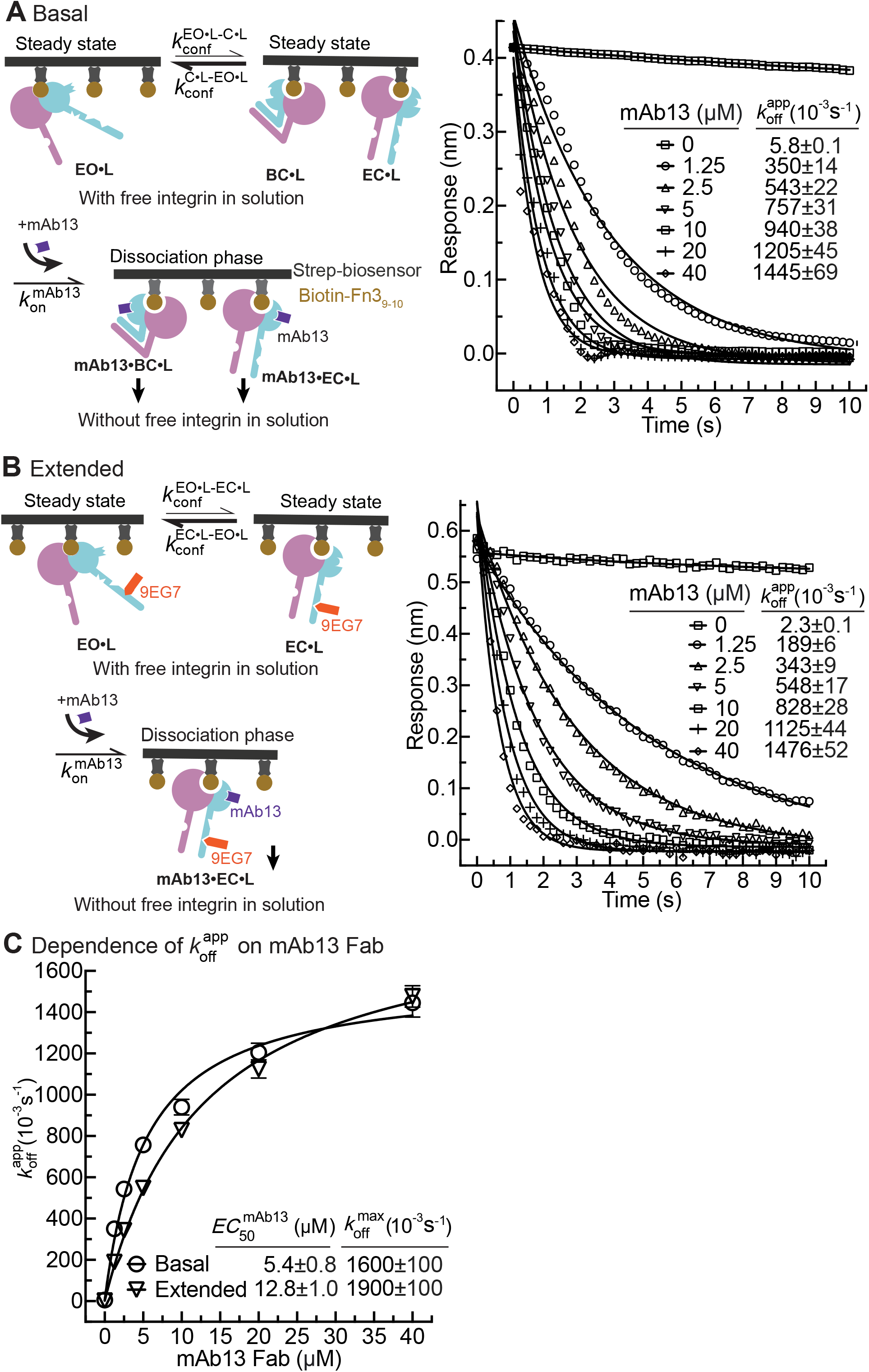
Dissociation of α5β1 ectodomain from biotin-Fn3_9-10_ in presence of closure-stabilizing Fab. (**A-B**) Unclasped high-mannose α5β1 ectodomain dissociation from biotin-Fn3_9-10_ immobilized on streptavidin biosensors was monitored by BLI. Reaction schemes are illustrated in each panel’s cartoons. Specifically, 50 nM α5β1 ectodomain was incubated with biotin-Fn3_9-10_ biosensors for 10 minutes to reach steady state binding in absence (A) or presence of 2μM 9EG7 Fab (B). Biosensors were then transferred into wells lacking the α5β1 ectodomain in presence or absence of 9EG7 Fab as before and also containing the indicated concentrations of mAb13 Fab for measurement of dissociation. The observed response (*R*_obs_) at each mAb13 Fab concentration as a function of time was individually fitted to the single exponential, 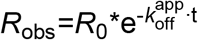, for the initial response at the start of dissociation (*R*_0_) and the 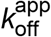. (**C**) Determination of 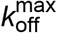 at saturating mAb13 Fab concentration. 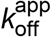 was fit to mAb13 Fab concentration using a dose response curve for the maximum off-rate at saturating mAb13 Fab concentration to determine 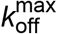. The mAb13 Fab concentration when the off-rate reaches half of the maximum, 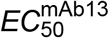 was also determined. Errors are from nonlinear least square fits.

### Calculation of ligand-binding kinetics from ensemble measurements

We directly measured the ligand-binding and dissociation kinetics for the EO state of α4β1 and α5β1 (Fig. 2C, E, Fig. 3B, Fig. 4C). In contrast, kinetics for the BC and EC states were only measured within ensembles. Their kinetics are convoluted in two respects. First, measurements on ensembles contain kinetics contributed by all states within the ensemble. Second, apparent association and dissociation kinetics may each contain a contribution from the kinetics of conformational change (Fig. 1B). Fig. 1B left shows apparent on- and off-rates and Fig. 1B right shows all the actual pathways by which ligand binding and dissociation can occur, which include all known integrin conformational states and the kinetics of conformational change between them. Furthermore, after ligand binding to the closed states, rapid conformational change to the EO state occurs and is responsible for our ability to measure the kinetics of binding as a result of accumulation of ligand-bound integrin in the EO state.

The underlying assumption for deconvoluting the kinetics of the closed states is that if integrin conformational transition kinetics are sufficiently fast so that the populations of the three integrin states do not deviate significantly during our experiments from the equilibrium values of the populations, then measured kinetics will not be significantly limited by conformational transition kinetics. In this case, both free integrins and ligand-bound integrins can be considered as readily equilibrated among their conformational states, and ligand binding coupled with integrin conformational changes can be approximated by the apparent 1 vs. 1 reaction between integrin and ligand (this allows the double tildes in Eqs. 1-4 in Fig. 7A to be treated as equal signs). All on- and off-rates measured here were well fit with the 1 vs. 1 Langmuir binding model (Fig. 2A-E, Fig. 3A-B, Fig. 4A-D, Fig 5B-C, and Fig. 6A-B), supporting this assumption. Moreover, reasonable agreement between the ratios of the apparent off- and on-rates, 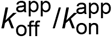, and previously determined equilibrium dissociation constants, *K*_d_, (Figs. 2F, 3C and 4F), validates the assumption that the apparent on- and off-rates (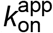 and 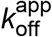) for each defined ensemble can be approximated by the on- and off-rates of each state weighted by its population in the ensemble (Fig. 7A, Eqs. 1-4). The population of the integrin states in absence of ligand (BC, EC, and EO) and in presence of saturating concentrations of ligand (BC•L, EC•L, and EO•L) were calculated based on the previously determined population and ligand-binding affinity of each state (Fig. S3B, Eqs. S5-S10) in the respective integrin α4β1 and α5β1 preparations (Li & Springer, 2018; Li *et al*., 2017) and are shown in Fig. 7B.

**Figure 7:**
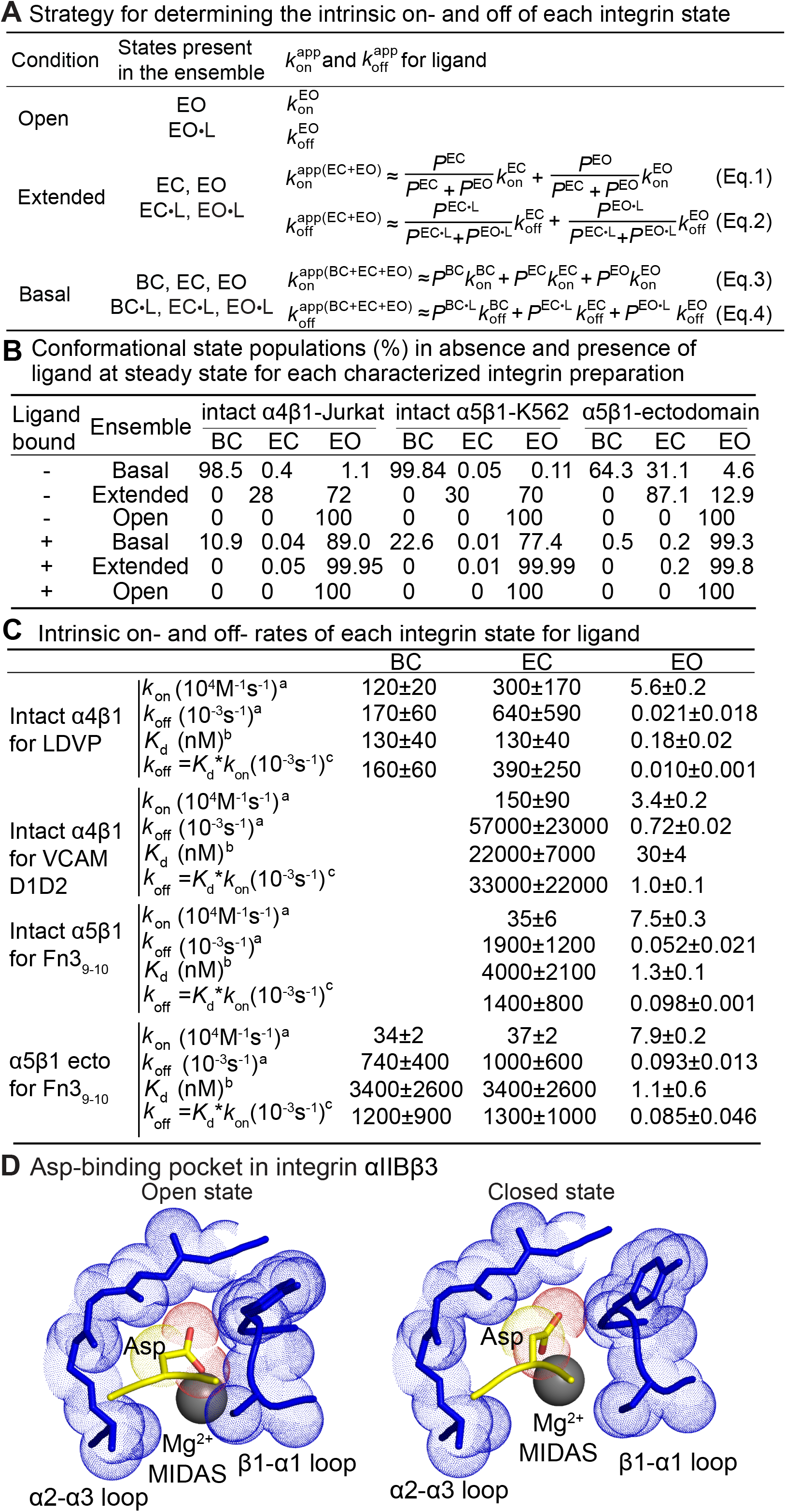
Ligand-binding kinetics of each integrin state. **(A)** Defined integrin α4β1 and α5β1 ensembles utilized in this study to measure ligand-interaction kinetics, as well as equations to relate the apparent on- and off-rates with the on- and off-rates for each conformational state. **(B)** Conformational state populations (%) in absence and presence of ligand at steady state. Previously reported populations for integrins in the absence of ligand and their affinities for ligand (Li & Springer, 2018; Li *et al*., 2017) were used with Eqs.S5-S10 in Fig. S3B to calculate the populations in saturating ligand of ligand-bound integrin states in each type of ensemble studied here. **(C)** Values of *k*_on_ and *k*_off_ for conformational states of four integrin-ligand pairs. As discussed in the text and Methods, kinetic measurements on the EO state and the extended and basal ensembles were used with equations in panel A to calculate kinetics of the BC and EC states. The errors for directly measured values were fitting errors from non-linear least square fit; the errors for calculated BC and EC values were propagated. ^a^: Intrinsic rates of EO state was from measurements in presence of HUTS4 & 9EG7 Fabs in Figs. 2-4, and intrinsic rates for BC and EC states were calculated with Eqs. 1-4 in panel A. ^b^: From equilibrium measurements as specified in Fig.2 to Fig.4 legends. ^c^: Calculated from the product of equilibrium *K*_d_ and *k*_on_. **(D)** Comparison of Asp-binding pocket in the open state (PDB: 3ze2 chains C+D) and closed state (PDB: 3zdy chains C+D) of integrin αIIBβ3 (Zhu *et al*., 2013). The pocket in the β3 βI domain is shown with backbone and nearby sidechains in blue stick and blue dot surfaces and the MIDAS Mg^2+^ ion as a silver sphere. The ligand Asp sidechain and its backbone loop are shown in yellow, red sidechain carboxyl oxygens. The Asp sidechain Cβ carbon and carboxyl oxygens are shown as yellow and red dot surfaces, respectively.

On- and off-rates for each α_4_β_1_ and α_5_β_1_ integrin state on intact cells and for the purified α5β1 ectodomain are summarized in Fig. 7C. Values are best determined, i.e. with the lowest errors, for the on-rate of EO state. Errors were higher for the BC and EC states, particularly for *k*_off_. Therefore, *k*_off_ values for each state were also calculated from *k*_off_ = *K*_d_\**k*_on_, where *K*_d_ is from equilibrium measurements (Li & Springer, 2018; Li *et al*., 2017). The *k*_off_ values of each state determined from these two strategies agree well with one another for each integrin-ligand pair.

In addition, *k*_off_ values were also comparable to the lower limit of 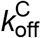 and 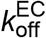 approached by measuring dissociation in presence of a closure-stabilizing Fab (Fig. 5 and Fig. 6).

## DISCUSSION

### Intrinsic ligand-binding kinetics of integrin conformational states

Employing conformation-specific Fabs against the integrin β1 subunit to stabilize integrin α4β1 and α5β1 into defined ensembles, we determined the on- and off-rates of each integrin conformational state. We found that despite ∼1,000 lower affinity, the closed states, BC and EC, of two β1 distinct integrins have markedly higher on-rates than the EO state. These findings have important implications for the sequence of events that occur when integrins interact with ligands, as discussed in the Integrin Activation section below. Previously, we determined equilibrium *K*_d_ values for the three different conformational states of integrins α4β1 and α5β1 (Li & Springer, 2018; Li *et al*., 2017). We used integrins on intact cells and as different types of ectodomain fragments. These different preparations differed up to 320-fold in affinity of their basal ensembles. However, integrin affinity for ligand was essentially identical for each integrin state and all differences in ensemble affinity were ascribable to variation among the preparations in the relative free energies of the three states. Therefore, we concluded that integrin affinity was intrinsic to each state (Li & Springer, 2018; Li *et al*., 2017). There may be real differences between cell surface and soluble integrins imposed by orientation, cell surface charge, and the glycocalyx; nonetheless, our previous measurements of *K*_d_ values for the EC and EO states of α5β1 on the cell surface and as an ectodomain fragment are within 2-fold of one another (Li & Springer, 2018; Li *et al*., 2017). These results are consistent with the intrinsic affinity concept, i.e. that integrin conformational state is the primary determinant of affinity, even though the geometry of integrins on cell surfaces may cause some modifications to these values that are minor compared to the large differences between the closed and open states.

Similar to intrinsic affinities, the results here on ligand-binding kinetics were consistent with on-rates and off-rates that are intrinsic to integrin conformational states. On-rates for intact α5β1 on cell surfaces and the α5β1 ectodomain in the EO state for the same fibronectin fragment were identical, and off-rates differed by only 1.8-fold. Similarly, on- and off-rates for the EC state of the intact cell-surface and ectodomain forms of α5β1 differed only by 1.1-fold and 1.9-fold respectively. We were able to measure on- and off-rates for the BC state of intact α4β1 binding to LDVP and for the BC state of the α5β1 ectodomain binding to Fn3_9-10_. In each case, the values of the BC state were within error of those for the EC state. The similar ligand-binding kinetics of the BC and EC states are in agreement with the essentially identical intrinsic affinities of the two closed states (Li & Springer, 2018; Li *et al*., 2017). In further agreement, crystal structures of the integrin αIIBβ3 ectodomain in the BC state and of the αIIBβ3 closed headpiece fragment, which has no interactions with the lower legs and thus serves as a model for the EC conformation (Zhu *et al*., 2008; Zhu *et al*, 2013), show essentially identical conformations of the ligand binding site.

We checked whether kinetics might be influenced by bound Fabs. In our previous work, we compared affinities measured with at least two Fabs specific for the closed, open, and extended states and for each state compared Fabs that bound to different domains. The results showed no significant differences between affinities measured with different Fabs. Here, we compared two Fabs used to stabilize the EO state and found slower association and dissociation kinetics with 12G10, which binds near the ligand binding site in the βI domain than HUTS4, which binds distally in the hybrid domain (Fig. 4F). As Fabs generally decrease dynamic protein motions in their epitopes (Wei *et al*, 2014) and may also sterically slow binding, the kinetics measured using HUTS4 Fab more likely approximate integrin kinetics in the absence of Fab and are repored in Fig. 7C.

The kinetics of the EC and BC states were calculated from measurements on extended or basal ensembles after correction for the kinetics in these ensembles contributed by the EO state. As a check on these measurements, we also measured *k*_off_ in the presence of mAb13 Fab, which after conformational conversion of EO•L to EC•L+BC•L trapped the closed states so that their dissociation could be measured. The lower limit of 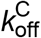 and 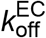 determined from these experiments (Fig.5C and 6C) are in good agreement with the calculated off-rate of the closed states (Fig. 7C).

Typical protein-protein on-rates as found for antibody-antigen interactions are in the range of 10^5^ to 10^6^ M^-1^ s^-1^ (Alsallaq & Zhou, 2008). The on-rates for the BC and EC states were in this range, e.g. 3.5×10^5^ and 1.5×10^6^ M^-1^ s^-1^ for α5β1 binding to Fn3_9-10_ and α4β1 binding to VCAM, respectively. In contrast, the on-rates for EO states for the corresponding integrin-ligand pairs were 7.5×10^4^ and 3.4×10^4^ M^-1^ s^-1^, respectively. These rates suggest a hindrance to ligand binding. Ligand-bound crystal structures in both open and closed conformations are known for two RGD-binding integrins, αIIBβ3 (Xiao *et al*, 2004; Zhu *et al*., 2013) and αVβ6 (Dong *et al*, 2014; Dong *et al*, 2017). Additionally, high resolution structures show RGD peptides bound to both closed and intermediate (partially open) conformations of α5β1 (Nagae *et al*, 2012; Xia & Springer, 2014). The open conformation has a tighter ligand-binding pocket. Slower ligand-binding kinetics for the open conformation is consistent with its tighter ligand-binding pocket, especially around the key RGD Asp reside (Fig. 7D). Movement of the β1-α1 loop toward the ligand and the MIDAS Mg^2+^ ion upon βI domain opening partially buries the Mg^2+^ ion and is expected to slow binding of the Asp sidechain, which must fit into a tight pocket with a specific geometry dictated by partially covalent and highly directional Asp sidechain metal coordination and hydrogen bonds to the β1-α1 loop backbone amide nitrogens.

The ∼1,000-fold higher affinities of the EO than the closed conformations for both α4β1 and α5β1 integrins are achieved by the ∼25,000-fold slower off-rate of the EO conformation (Fig. 7C). Similar to the differences in on-rates, the differences in off-rates can be understood in terms of the structural details in the ligand-binding pocket and the much higher affinity of the EO state. The tighter Asp sidechain binding pocket and greater burial of the Asp provide a barrier to dissociation (Fig. 7D). The number of hydrogen bonds of the Asp sidechain to the β1-α1 loop backbone increases in the open state (Zhu *et al*., 2013). Furthermore, the greater burial of these polar bonds and increased network of hydrogen bonds around them increases their strength.

The Arg sidechain also strengthens its hydrogen bonding in the open state. During opening, as the βI domain β1-α1 loop moves toward the Asp, the entire RGD moiety slides toward the α-subunit, which is to the left in the view of Fig. 7D. This movement is seen in Fig. 7D as the closer approach of the Asp to the α2-α3 loop in the open state. A hydrogen bond network with two waters between the RGD Arg sidechain and α5β1 residue Gln-221 in the closed state is exchanged for a direct Arg hydrogen bond to α5β1 Gln-221 as RGD slides toward α5β1 during opening (Xia & Springer, 2014). All these can contribute to the higher affinity and tens of thousands-fold slower off-rate of the EO conformation than the closed conformations.

Our kinetic measurements were carried out at 22°C. At 37°C, both on- and off-rates will be higher. Increase in temperature generally has a much greater effect on dissociation rates than association rates (Johnstone *et al*, 1990). The amount of increase depends on the activation energy; i.e. the height of the energy barrier to dissociation.

The intrinsic ligand-binding kinetics of integrin conformational states described here are consistent with previous kinetic observations. These studies showed that activating integrin ensembles with Mn^2+^ or activating IgG or Fab, using conditions that in retrospect would partially, but not completely, shift integrin ensembles to the EO state, decreased the ligand off-rates of integrins α4β1 and α5β1 (Chigaev *et al*, 2001; Takagi *et al*., 2003). The extremely long lifetime of the α5β1 complex with fibronectin in the EO state, around several hours, explains why the α5β1 complex with fibronectin in Mn^2+^ was much more rapidly reversed by mAb 13 IgG specific for the closed conformations than by competitive inhibitor (Mould *et al*, 2016; Mould *et al*., 2014).

### Integrin activation

A major impetus for these studies was to determine the pathway for activation of integrins in cells, i.e. the activation trajectory. Of key importance is how integrins on the cell surface first engage ligands. By dynamically linking the actin cytoskeleton to the extracellular environment, integrins transduce both external and internal mechanochemical cues and bi-directionally signal across the plasma membrane. Integrin signaling is governed by cytoskeletal force and the force stabilized, high-affinity, extended-open conformation is the only state competent to mediate cell adhesion (Alon & Dustin, 2007; Astrof *et al*., 2006; Li & Springer, 2018; Li *et al*., 2017; Nordenfelt *et al*., 2016; Nordenfelt *et al*., 2017; Sun *et al*., 2019; Zhu *et al*., 2008). Mechanotransduction occurs when integrins bind to ligand anchored in the extracellular environment, the cytoplasmic domain simultaneously binds to a cytoskeletal adaptor and links to actin retrograde flow, and a tensile force is transmitted through the integrin that stabilizes the extended-open conformation over the bent-closed conformation. Thus the on- and off-rates of ligand binding to integrins are among the key parameters that determine the cytoskeletal force regulation efficiency. We found that the closed states, with loose ligand binding pockets, have higher on-rates for ligand binding, making them the most efficient state for encountering ligand.

Because the BC state is >200-fold more populated than the EC state for both integrins α4β1 and α5β1 on the cell surface (Li & Springer, 2018; Li *et al*., 2017) (Fig. 7B), the BC state may have an important role in initial binding to ligand. There are few constraints on the orientation of integrins on cell surfaces until transmitted force orients them when they bridge extracellular ligands and the actin cytoskeleton (Nordenfelt *et al*., 2017; Swaminathan *et al*, 2017). In the absence of such engagement, linkers of largely disordered residues between the last module of integrin α- and β-subunit ectodomains and the beginning of their transmembrane α-helices allows large tilting motions of the ectodomain relative to the plasma membrane (Zhu *et al*, 2009). Thus, the common depiction of the leg domains of integrins and other receptors as oriented normal to the cell membrane (Fig. 1A) is only a conventional cartoon representation and has no experimental basis. Structures of integrin αIIBβ3 linker and transmembrane domains on cell surfaces, combined with the ectodomain, showed that large movements of the BC state relative to the membrane normal were possible. Nonetheless, none of these orientations have a ligand binding site with an orientation optimal for binding a ligand on the surface of another cell or in the extracellular matrix. In contrast, the ligand binding site is better exposed in the EC state (Fig. 1A) and the EC state also can bend at multiple domain-domain junctions and is less constrained in orientation relative to the plasma membrane. It is possible that either the BC state is the predominant ligand-binding state, or that the BC state provides a large reserve of integrins that, through conformational sampling of the EC state, allows the EC state to be the predominant ligand-binding state.

Once ligand is bound to the BC or EC state, the ∼1000-fold higher ligand-binding affinity for the EO conformation strongly favors conformational change to the EO state (Li & Springer, 2018; Li *et al*., 2017). If an adaptor and the actin cytoskeleton are bound at the time when a ligand that is embedded in the extracellular environment is bound to the integrin, the ligand resists the force from actin retrograde flow, and tensile force is transmitted through the integrin, strongly stabilizing the EC•L and EO•L conformations (Li & Springer, 2017). Furthermore, the EO•L state is ∼2,000 and ∼10,000 more populated than the EC•L state for integrins α4β1 and α5β1, respectively (Fig. 7B). The off-rates of the EO states of α4β1 and α5β1 equate to lifetimes of about 0.4 hours and 3 hours, respectively, and serve to make integrin-ligand bonds highly resistant to detachment. In contrast, when engagement to the adaptor/actin cytoskeleton is reversed, BC•L would become substantially populated in the basal ligand-bound conformations (Fig. 7B), and with a lifetime in millisecond to second range, would allow the integrin to dissociate from ligand.

In summary, we have substantially advanced our understanding of how integrins on intact cells bind ligands by measuring the ligand binding and dissociation kinetics for the three conformational states of two integrins, α4β1 and α5β1. While it may seem surprising that the low affinity states bind more rapidly than the high affinity states, our findings concord with previous studies on selectins (Phan *et al*., 2006) and bacterial fimbriae adhesins (Yakovenko, 2015) that have two states, one flexed (bent) and the other extended, that are also subjected to regulation by force, in which the extended state is the higher affinity state. As there is no structural homology between the three classes of adhesins, convergent evolution appears to have selected a low affinity, flexed/bent state for rapid ligand binding that can be subsequently stabilized by force to a high affinity, extended state that can then better resist the tendency of force to accelerate receptor-ligand dissociation. In the two-state systems, force is applied externally by shear flow, while in the three-state integrin system, force is applied internally by engagement of actin retrograde flow. This empowers the actin cytoskeleton machinery to regulate integrin function, ensuring intimate coordination between the needs of adhering and migrating cells because the same signaling pathways that regulate actin polymerization and disassembly also regulate formation of cellular attachments through integrins to the extracellular environment. While the kinetics of integrin conformational change remain to be measured, the excellent fit of kinetic measurements to the 1 vs. 1 Langmuir model found here and agreement between *k*_off_/*k*_on_values and *K*_d_ measured at equilibrium suggest that integrin conformational change kinetics are also rapid. Rapid ligand binding, together with rapid cytoskeletal adaptor binding, would enable their coincidence to regulate integrin activation, thus providing a seamless method for activating integrins at cellular locations where actin is activated and at extracellular locations where ligand is available.

## MATERIALS AND METHODS

### Fabs

IgGs, 8E3 (Mould *et al*, 2005), 9EG7 (Bazzoni *et al*, 1995), 12G10 (Mould *et al*, 1995), HUTS4 (Luque *et al*, 1996), mAb13 (Akiyama *et al*, 1989) and SNAKA51 (Clark *et al*, 2005) were produced from hybridomas and purified by protein G affinity; Fabs were prepared with papain digestion in PBS (phosphate-buffered saline with 137 mM NaCl, 2.7 mM KCl, 10 mM Na2HPO4 and 1.8 mM KH2PO4, pH7.4) with 10 mM EDTA and 10 mM cysteine and papain: IgG mass ratio of 1:500 for 8 hrs at 37°C, followed by Hi-Trap Q chromatography in Tris-HCl pH 9 with a gradient in the same buffer to 0.5 M NaCl.

### Integrin α5β1 soluble preparations

Integrin α5β1 ectodomain (α5 F1 to Y954 and β1 Q1 to D708) with secretion peptide, purification tags, and C-terminal clasp (Takagi *et al*, 2001) were produced by co-transfecting the pcDNA3.1/Hygro(-) vector coding the α-subunit and pIRES vector coding the β-subunit into HEK 293S GnTI^-/-^(*N*-acetylglucosaminyl transferase I deficient) cells. Stable transfectants were selected with hygromycin (100 μg/ml) and G418 (1 mg/ml), and proteins were purified from culture supernatants by His tag affinity chromatography and Superdex S200 gel filtration after cleavage of C-terminal clasp and purification tags with Tev protease (Li *et al*., 2017).

### Peptidomimetic and macromolecule fragments

FITC-conjugated α4β1 specific probe, 4-((N′-2-methylphenyl)ureido)-phenylacetyl-L-leucyl-L-aspartyl-L-valyl-L-prolyl-L-alanyl-L-alanyl-L-lysine (FITC-LDVP) and its unlabeled version, LDVP, were from Tocris Bioscience (Avonmouth, Bristol, United Kingdom). Human VCAM D1D2 (mature residues F1 to T202) were expressed and purified from HEK 293S GnTI^-/-^cell line supernatants by affinity chromatography and gel filtration (Yu *et al*, 2013). VCAM D1D2 was fluorescently labeled with Alexa Fluor 488 NHS Ester (ThermoFisher Scientific). Human Fn3_9-10_ S1417C mutant (mature residues G1326 to T1509) and its synergy and RGD sites (R1374A&P1376A&R1379A&S1417C&Δ1493-1496) mutated inactive version were expressed in *E. coli* and purified as described (Li *et al*., 2017; Takagi *et al*., 2001). Fn3_9-10_ S1417C mutant was fluorescently labeled with Alexa Fluor 488 C5 maleimide (ThermoFisher Scientific) at residue Cys-1417. Both Fn3_9-10_ S1417C mutant and its inactive version were biotinylated with Maleimide-PEG11-Biotin at residue 1417 (ThermoFisher Scientific) in PBS.

### Quantitative fluorescent flow cytometry

Jurkat and K562 cells (10^6^ cells/mL in RPMI-1640 medium, 10% FBS) were washed twice with assay medium (Leibovitz’s L-15 medium, 10 mg/mL BSA) containing 5 mM EDTA, twice with assay medium alone, and suspended in assay medium. Cells at 2×10^6^ cells/mL were incubated with indicated concentration of Fabs for 30min at 22°C. Addition of FITC-LDVP, Alexa488-VCAM D1D2 (1.6 labeling ratio) or Alexa488-Fn3_9-10_ (1.0 labeling ratio) at indicated concentrations initiated association. Association was measured as mean fluorescence intensity (MFI) at successive time points after addition of the fluorescent ligands. Addition of 500-fold higher concentration of the unlabeled ligand at the end of the association phase initiated the dissociation phase. Background MFI for FITC-LDVP, Alexa488-VCAM D1D2 and Alexa488-Fn3_9-10_ in presence of 10 mM EDTA was subtracted (Supplemental Fig. S1).

### Fitting flow cytometry and BLI kinetic binding traces with 1 vs. 1 Langmuir binding model

Kinetic traces including both the association phase and the dissociation phase at different analyte concentrations were globally fitted to the following function.

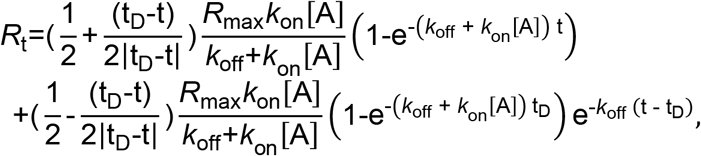

where t is time, *R*_t_ is response at time t, t_D_ is the time that dissociation starts, [A] is the analyte concentration, and *R*_max_ is the maximum response. The first term fits the data in the association phase and the second term fits the data in the dissociation phase. The prefactor of the first term is 1 prior to t_D_ and becomes 0 after t_D_; whereas the prefactor for the second term is 0 prior to t_D_ and becomes 1 after t_D_. Nonlinear least square fit of *R*_t_, [A], and t to the above equation yields the on-rate, *k*_on_, off-rate, *k*_off_, and *R*_max_.

### Bio-Layer Interferometry (BLI)

Binding kinetics of unclasped high-mannose α5β1 ectodomain and Fn3_9-10_ was measured by BLI (Wallner *et al*., 2013) with streptavidin biosensors on an Octet RED384 System. The reaction was measured on 96 well plate (200 uL/well) in buffer with 20 mM Tris HCl (pH 7.4), 150mM NaCl, 1mM Ca^2+^, 1mM Mg^2+^ and 0.02% Tween20. Streptavidin biosensors were hydrated in reaction buffer for 10 min before starting the measurements. Each biosensor was sequentially moved through 5 wells with different components: (1) buffer for 3 minutes in baseline equilibration step; (2) 35 nM biotin-Fn3_9-10_ for 1 minute for immobilization of ligand onto the biosensor; (3) indicated concentrations of Fabs for 5 minutes for another baseline equilibration; (4) indicated concentrations of α5β1 ectodomain and Fabs for the association phase measurement; (5) indicated concentrations of Fabs for the dissociation phase measurement. Each biosensor has a corresponding reference sensor that went through the same 5 steps, except in step 2 the ligand was replaced with 35 nM inactive version of Fn3_9-10_ with both the RGD binding site and the synergy site (PHSRN) mutated. Background subtracted response in both the association and dissociation phases, and at different α5β1 ectodomain concentrations, were globally fit to the 1 vs. 1 Langmuir binding model, with 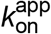 and 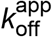 as shared fitting parameters and maximum response (*R*_max_) for each biosensor as individual fitting parameter. The equilibrium binding (response) was calculated from 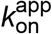 and 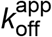 values at each each α5β1 ectodomain concentration and fit to a dose response curve to calculate *K*_d_ values as a check on 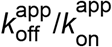 values. To calculate equilibrium response (*R*_eq_), fitted *k*_on_, *k*_off_, and *R*_max_ values at each α5β1 ectodomain concentration [A] were used to calculate *R*_eq_ at a time 1,000-folder longer than the “binding time”, i.e., 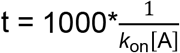 with the following equation:

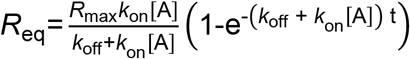

### Calculating ligand-binding and dissociation rates for the BC and EC states

The measured on- and off-rates (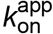 and 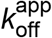) for each defined ensemble containing 2 or 3 states shown in Figs. 2-4 was approximated by the on- and off-rates of each state weighted by its population in the ensemble (Fig. 7A, Eqs. 1-4). At steady state, the population of the free integrin states and the ligand-bound integrin states were calculated based on the previously determined population and intrinsic ligand-binding affinity of each state (Fig. S3B, Eqs. S5-S10) in the respective integrin α4β1 and α5β1 preparations (Li & Springer, 2018; Li *et al*., 2017) (Fig. 7B). Specifically, 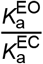 and 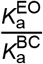 (in Fig.S3B, Eqs.S8-S10), are the intrinsic ligand-binding affinity ratios of the EO state and the closed states. For integrin α4β1, the ratios were averaged to 745±237 from six α4β1 preparations, including α4β1 headpiece with high-mannose N-glycans, α4β1 ectodomain with high-mannose N-glycans, α4β1 ectodomain with complex N-glycans, and intact α4β1 on three different cell lines (Li & Springer, 2018); for integrin α5β1, the intrinsic ligand-binding affinity ratio of the EO state and the closed states were averaged to 3106 ±1689 from eight soluble α5β1 preparations that varied in presence or absence of the lower legs, of a loose clasp in place of the TM domain, and in whether the N-linked glycan was complex, high mannose, or shaved (Li *et al*., 2017). Using 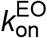 and 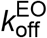 rates experimentally measured in Figs. 2-4, 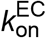 and 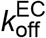 were derived from the 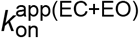 and 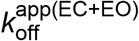 measured in extended ensembles, respectively (Fig. 7A, Eqs.1-2). By including the values for 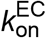 and 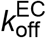 in addition to 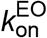 and 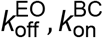 and 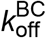 were then derived from 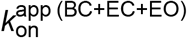 and 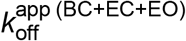 measured in basal ensembles, respectively (Fig. 7A, Eqs. 3-4).

## Acknowledgements

We thank Kelly L. Arnett in Center for Macromolecular Interactions of Harvard Medical school for training and consultation on BLI measurement. We thank Taekjip Ha for suggestions on our manuscript. This work was funded by NIH R01 HL131729 (“Activation trajectories of integrin a5b1”).

## Author Contributions

J.L. and T.A.S. designed research and wrote the paper. J.L. carried out the measurements and analyzed the data. J.B.Y. prepared the Fabs and purified soluble integrins and ligands.

## Conflict of Interest Statement

The authors declare no competing financial interests.

